# The vectoring competence of the mite *Varroa destructor* for Deformed wing virus of honey bees is highly dynamic and affects survival of the mite

**DOI:** 10.1101/2022.05.06.490899

**Authors:** Eugene V. Ryabov, Francisco Posada-Florez, Curtis Rogers, Zachary S. Lamas, Jay D. Evans, Yanping Chen, Steven C. Cook

## Abstract

The ectoparasitic mite, *Varroa destructor* and the viruses it vectors, including types A and B of Deformed wing virus (DWV), pose a major threat to honey bees, *Apis mellifera*. Analysis of 256 mites collected from the same set of colonies on five occasions from May to October 2021 showed that less than a half of them, 39.8% (95% confidence interval (CI): 34.0 - 46.0%), were able to induce an overt-level DWV infection with more than 10^9^ viral genomes per bee in the pupa after 6 days of feeding, with both DWV-A and DWV-B being vectored at similar rates. To investigate the effect of the phoretic stage on the mites’ ability to vector DWV, the mites from two field collection events were divided into two groups, one of which was tested immediately for their infectiveness, and the other was kept with adult worker bees in cages for 12 days prior to testing their infectiveness. We found that while 39.2 % (95% CI: 30.0 – 49.1%) of the immediately tested mites induced overt-level DWV infections, 12-day phoretic passage significantly increased the infectiousness of mites to 89.8% (95% CI: 79.2 – 95.6%). It is likely that Varroa mites that survive brood interruptions in field colonies are increasingly infectious. We found that mite lifespan was significantly affected by the type of DWV it transmitted to pupae. The mites, which induced overt DWV-B but not DWV-A infection had an average lifespan of 15.5 days (95% CI: 11.8 - 19.2 days), which was significantly shorter than those of the mites which induced overt DWV-A but not DWV-B infection, with an average lifespan of 24.3 days (95% CI: 20.2 - 28.5), or the mites which did not induce high levels of DWV-A or DWV-B, with an average survival of 21.2 days (95% CI: 19.0 - 23.5 days). The mites which transmitted high levels of both DWV-A and DWV-B had an intermediate average survival of 20.5 days (95% CI: 15.1 - 25.9 days). The negative impact of DWV-B on mite survival could be a consequence of the ability of DWV-B, but not DWV-A to replicate in Varroa mites.

## Introduction

Survival of pathogens is dependent on their ability to infect new host individuals. Although vertical transmission plays a role in maintaining pathogens in host populations, it is horizontal transmission that allows pathogens to be maintained in host populations and to extend their host range (1). It is also widely accepted that horizontal transmission, which does not rely on long-term host survival and reproductive success, favors the emergence and existence of highly virulent variants of pathogens (1–3). Routes of horizontal transmission may include fecal-oral, respiratory and mechanical, as well as transmission by other living organisms, *i.e*. disease vectors (4). Vector-mediated transmission greatly increases pathogens’ spread because it allows their movement between host individuals separated spatially or temporarily, allows vectors to act as a reservoir for pathogens, and delivers pathogens directly into susceptible host tissues and cells, thereby bypassing protective barrier tissues, as in the cases of hematophagous arthropod vectors feeding on vertebrates’ blood or aphids feeding on plant phloem (5). Vector-mediated transmission either could and/or could not involve replication of the pathogen. In the case of replicative transmission, the requirement to be adapted to both host(s) and vector affects the pathogen evolution.

Arthropods, such as insects and ticks, are common and well-studied vectors of viral pathogens of vertebrates and plants, but their vectoring of viruses that infect insects is far less common and poorly characterized. An important example of this latter case includes the vectoring by the ectoparasitic mite, *Varroa destructor* (Varroa), of viruses that infect the Western honey bee, *Apis mellifera*, a principal managed insect pollinator. Over the last decades, honey bees in Europe and North America have suffered increased colony losses caused by the spread of Varroa, which moved from its original host, the Asian bee, *Apis cerana*, in the 1900s (6), and has quickly become nearly cosmopolitan in distribution. Varroa feed on hemolymph and fat body of honey bee adults and pupae (7, 8) and can transmit a number of viruses, including Deformed wing virus (DWV) type A (DWV-A) (9) and type B (DWV-B) (10). DWV was present in pre-Varroa honey bees (11), and is present in honey bees in Varroa-free regions of the globe, but infections are mainly asymptomatic and the virus accumulates to low levels. Mite-mediated vectoring provides an efficient route of horizontal transmission for both variants of DWV, DWV-A and DWV-B (12), leading to selection of highly virulent variants of DWV, which can reach high levels in honey bees (13–15).

Recently it was shown that DWV-B genomic RNA was present in Varroa epithelial cells while DWV-A genomic RNA was not, strongly suggesting that DWV-B is replicating in mites (16). The lack of replication of DWV-A in Varroa could explain previously reported non-replicative, and possibly even non-persistent transmission of DWV-A by the mite (17). Moreover, passaging field-collected mites on virus-free pupae significantly reduced the mites’ ability to transmit DWV-A, suggesting that DWV-A vectoring is non-persistent (17). The ability of DWV-B to replicate in its mite vector may lead to the virus persistence in Varroa resulting in more efficient vectoring compared to DWV-A. At the same time, replication of DWV-B in Varroa may have a direct impact on mite fitness.

Here, we studied vectoring of DWV variants A and B by Varroa in controlled laboratory experiments to determine (1) the competence of phoretic and pupae-associated mites from field colonies to vector DWV to naïve pupae, (2) how a prolonged phoretic stage on adult bees impacts the vectoring competence of Varroa, and (3) whether the transmission of the different DWV strains impacts Varroa survival.

## Materials and Methods

### Varroa mites and honey bees

Experiments to determine Varroa vectoring competence were carried out between May to October 2021 at the USDA Bee Research Laboratory, Beltsville, Maryland with both Varroa mites and honey bees collected from honey bee colonies housed at the laboratory apiary.

Varroa were sourced from an isolated group of 4 colonies located at 39°02’26”N 76°51’41”W, which did not receive varroacide treatment during 2020 and 2021. These colonies showed approximately 3% mite infestation rate in May, increasing to 5%in October. Phoretic mites were sourced from adult bees collected from Varroa-infested hives using the sugar roll method (18), while pupa-associated adult mites were collected from the capped brood cells in the same colonies. Mites from all four colonies of the Varroa-infested apiary were pooled to obtain enough mites for experiments. To avoid starving mites, within one hour of collection, mites were placed on bee hosts (pupae the vectoring competence experiments or worker bees for the cage experiments (**Figure 1**). Mites were inspected by microscope and only actively moving individuals were used.

**Figure 1.**
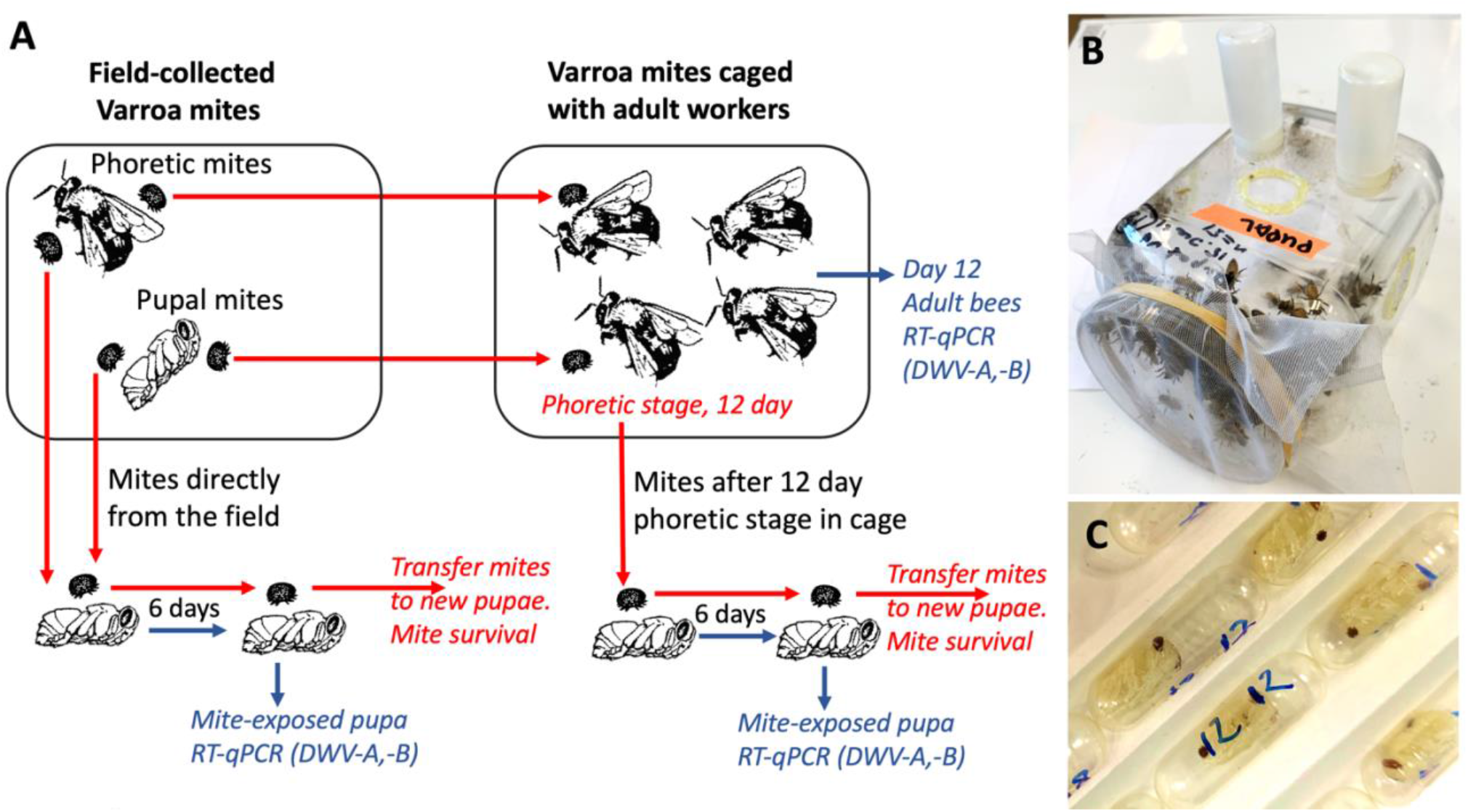
A. Schematic representation of the experiments. Varroa mite vectoring competence for DWV was tested by exposing naïve honey bee pupae to individual mites in gelatin capsules. Mites were either placed immediately after collection from filed colonies (field-collected) or were passaged on adult worker bees in cages. Mite survival on the pupae was recorded. B. Cage with adult worker honey bees. C. Gelatin capsules with mites feeding on the honey bee pupae.

Recipient pupae and newly emerged worker bees were obtained from two honey bee colonies located at 39°02’32”N 76°51’53”W, which had received varroacide treatment in the Autumn 2020. The source colonies had relatively low mite infestation rates, approximately 0.5 % in May and June, and approximately 3% in August to October. The colonies which were used to source mites and pupae were monitored for the presence of common honey bee viruses, and were shown to be free of Sacbrood virus, Acute bee paralysis virus, Israeli acute paralysis virus, Kashmir bee virus, Black queen cell virus, Chronic bee paralysis virus, and Lake Sinai virus, while both DWV-A and DWV-B were detected. The white-eye stage pupae destined for the mite virus vectoring competence experiments (**Figure 1A**) were pulled out from the brood cells using soft tweezers, and both pupae and their brood cells were inspected to exclude pupae that were previously exposed to Varroa mite feeding and could be already infected with viruses. To obtain worker bees for the cage experiments (**Figure 1B**), frames of capped brood were placed in an incubator set at +33°C, relative humidity (RH) 85%. After 18 hours and the newly emerged adult worker bees were collected for experiments.

### Adult bee cage experiments

To investigate effects of a prolonged phoretic stage on the mites’ vectoring competence for DWV-A and DWV-B (*i.e*. ability to transmit these viruses to bees), mites collected from honey bee colonies were kept on newly emerged adult worker bees in laboratory cages for 12 days prior to pupal infectiveness test. Groups of 150 to 161 newly emerged worker bees from the colonies with low Varroa counts were housed in ventilated, sterilized plastic cages measuring 15 cm by 20 cm by 20 cm and maintained in a dark incubator set at 33.0 °C and having 85% RH. The bees were given *ad libitum* access to both 1: 1 sugar syrup and tap water contained in separate feeders, which were each comprised of an inverted 20 mL plastic vial with holes drilled into their caps. Both syrup and water feeders were changed every 24 h. Field-collected Varroa were added to the cages at the same day as worker bees. Separate cages were used for the phoretic and pupae-associated mites and for a control cage group which did not contain mites. Dead bees and mites were counted and removed daily. After 12 days, all surviving mites were collected from remaining bees in the cages, and then individually transferred from the worker bees to separate naïve pupae in gelatin capsules for vectoring competence tests (**Figure 1**), see below. Random samples from the remaining live adult bees were collected and immediately frozen at −80°C. Total RNA was extracted from frozen individual workers, and then used to quantify DWV-A and DWV-B copy numbers using the reverse transcription-quantitative polymerase chain reaction (RT-qPCR). The oligonucleotide primers (5’-GAGATTGAAGCGCATGAACA-3’ and 5’-TGAATTCAGTGTCGCCCATA-3’) and 3’-CTGTAGTTAAGCGGTTATTAGAA-3’ and 5’-GGTGCTTCTGGAATAGCGGAA-3’) were used for quantification of DWV-A and DWV-B, respectively. A dilution series of the recombinant plasmid containing full-length cDNA of DWV-A (19) and DWV-B (20) ranging from 1 to 10^9^ genomic copies was used to establish a standard curve for by plotting Ct values against the log-transformed cDNA concentrations. The results of DWV-A and DWV-B quantifications are summarized in Supplementary Tables S1-S4.

### Analysis of Varroa virus vectoring competence and survival

Varroa virus vectoring competence tests were similar to those described in (17) and involved placing individual mites, either directly collected from field colonies or after a 12-day period phoretic stage on adult bees in cages, on naïve white-eye honey bee pupa in a gelatin capsule for 6 days in a dark incubator set at +33.0 °C and having 85% RH to allow mite feeding and thereby assess virus infection of these pupae. For each infectivity test experiment, a set of control pupae from the same collection was incubated in gelatin capsules without Varroa. Mite survival was monitored every 24 hours. After six days, all mite-exposed pupae were removed and stored at −80°C. Total RNA preparations were extracted from frozen pupae and were used to quantify viral copy numbers via RT-qPCR as reported in (20). The mites which survived incubation on pupae were transferred to fresh naïve white-eye pupae in the same capsules and daily mite survival monitoring continued. Every 6 days, surviving mites were transferred to new naïve pupae. Mite survival data are summarized in **Supplementary Table S5**.

### Statistical analysis

R statistical package was used for statistical analysis (21). Log-transformed virus levels were used for paired t-tests. Confidence intervals were calculated by the modified Wald method (22). The nonparametric Wilcoxon signed-rank test was used for the mite survival analysis.

## Results

### Infectivity tests showed that a significant proportion of field-collected Varroa mites did not vector DWV

The ability of field-collected Varroa mites to infect honey bee pupae with DWV-A or DWV-B was investigated at five timepoints from May to October to determine if changes of vectoring competence occur seasonally. In total, the study involved 256 mites, both phoretic, from adult bees (n=143), and pupal, from capped brood cells (n=113). It was found that for all mites from the five collections, the average level of DWV-A in pupae exposed to phoretic mites (7.81 ± 2.01 SD log_10_ genome equivalents (GE)/ pupae, mean ± SD), and in the pupal mite-exposed recipient pupae (8.24 ± 2.07 GE/pupae, mean ± SD), were significantly higher than that in the control mite non-exposed pupae (6.50 ± 1.00 log_10_ GE/pupa, mean± SD), pairwise t-test, *P* < 0.001, **Figure 2A**), but for the pupae of both Varroa-exposed groups the virus levels were not significantly different, pairwise t-test, P = 0.070 (**Figure 2A**). Similar vectoring competence was observed for DWV-B, while there were no significant differences of DWV-B levels between pupae exposed to phoretic and pupal mites (7.49 ± 2.20 log_10_ GE/pupa and 7.93 ± 2.33 2.20 log_10_ GE/pupa, correspondingly, *P* = 0.097), these levels were significantly higher than in control pupae (6.70 ± 1.07 log_10_ GE/pupa; pairwise t-test, *P* < 0.05; **Figure 2B**).

**Figure 2.**
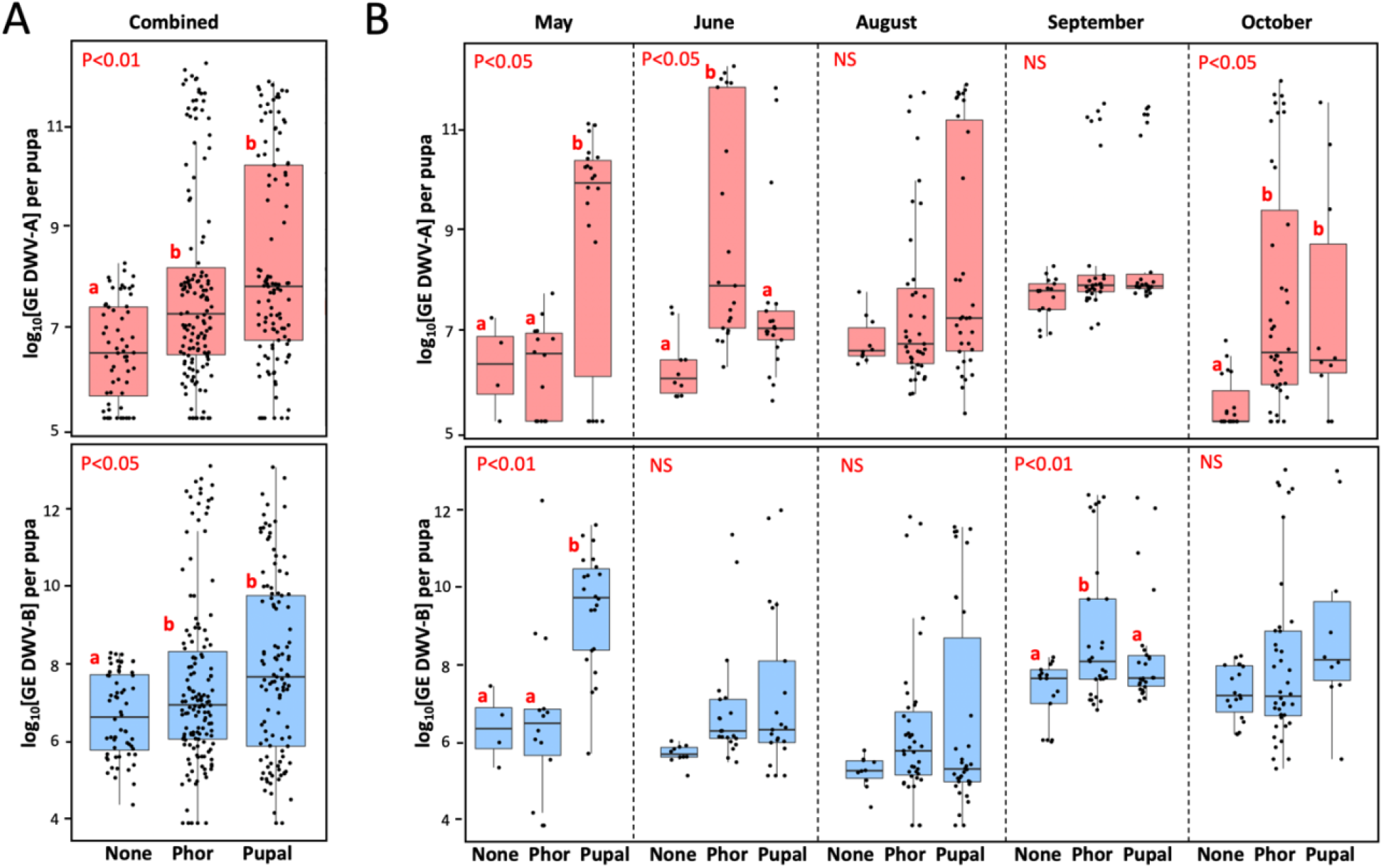
Vectoring competence of field-collected Varroa mites for DWV-A (upper panels) and DWV-B (lower panels) for five collection events combined (A) and for individual collection events (B). Boxplots show development of virus infection in individual naïve pupae after 6-day exposure with the phoretic, “Phor”, or pupal, “Pupa”, mites. Non-exposed control pupae - “None”. Dots represent virus levels in individual pupae. Red letters above boxes indicate significantly and non-significantly different groups (pairwise t-test), NS – non-significant. Separate statistical analyses were carried for each collection event.

Importantly, we found that none of the pupae not exposed to mites (n=60) showed levels of either DWV-A or DWV-B exceeding 10^9^ GE (0.0%, CI 95%: 0.0 - 7.2%). In contrast, 36.4% (CI 95%: 28.9 - 45.5%) of recipient pupae exposed to mites had developed overt level infections of either DWV-A or DWV-B. Similar vectoring capacities were observed for DWV-A and DWV-B for both phoretic mites (22.4%, CI 95%: 16.3 - 29.9 %, and 17.5%, CI 95%: 12.1 - 24.6 %) and pupal mites (33.6 %, CI 95%: 25.6 - 42.8 %, and 31.9%, CI 95%: 24.0 - 40.9 %), **Figure 2A**.

Infectiveness of both phoretic and pupal mites varied considerably between five collection events from May to October. In some collections, no significant difference between average DWV-A or DWV-B was observed between the control pupae and the mite-exposed pupae (**Figure 2B**). Notably, in every collection event there were mites which induced overt DWV infection (>10^9^ GE) in the recipient pupae (**Figure 2B**, “Phor”, “Pupal”), while no overt DWV infection developed in the control pupae (**Figure 2B**, “None”). Although we observed a significant variation of infectiveness of the pupal and phoretic mites collected in different months, we observed no seasonal trend in mite vectoring competence for DWV from May to October (**Figure 2B**). Since levels of DWV-A and DWV-B in naïve pupae which were not exposed to mites varied through the season, statistical analysis was carried out for each collection separately (**Figure 2B**).

### Vectoring competence of Varroa for DWV increased following a phoretic stage

Varroa reproduces exclusively in capped brood cells (23), therefore brood interruption can be used to reduce Varroa loads (24). During broodless periods, mites survive a prolonged phoretic stage on adult bees [25], but little is known how such phoretic stage impacts on the ability of Varroa to vector viruses. To test this experimentally, we used phoretic and pupal mites from the June and September field collection events. The mites were randomly divided into two groups, one of which was tested immediately after collection for their vectoring competence for DWV-A and DWV-B on the naïve pupae (**Figure 3**, “Field-collected”), and another group was kept on newly emerged adult worker bees as phoretic in laboratory cages for 12 days before the Varroa vectoring competence tests. In both June and September experiments, worker bees suffered considerable losses by day 12. In the June cage experiment, where there were 150 worker bees in each of three cages, the numbers of surviving bees by day 12 were 92 for the no-mite control group, 56 for the phoretic mite-exposed and 17 for pupal-mite exposed groups. In the September experiment, in which 150, 161 and 155 worker bees were placed in control, “Phor”, “Pupal” cages, respectively, after 12 days 120, 96, and 15 bees survived in the respective cages. In both experiments, losses of honey bee workers in the cages infested with mites were significantly higher than in the control mite-free cages (*P* < 0.0001, Pearson’s chi-square test). Also, the losses of worker bees were significantly higher in the cages with the “Pupal” mites compared with the “Phor” mites (*P* < 0.0001, Pearson’s chi-square test).

**Figure 3.**
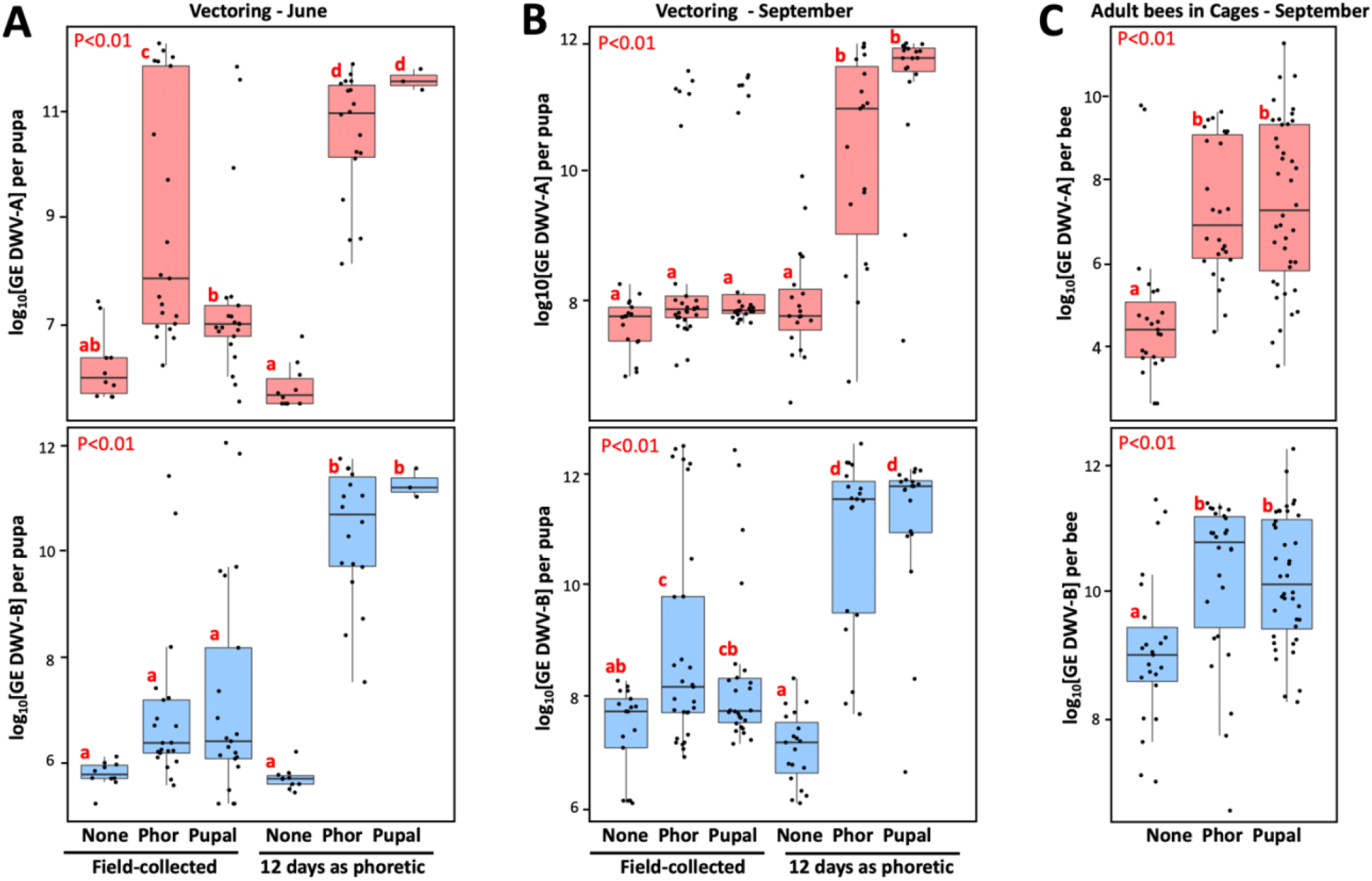
Effect of phoretic passage on Varroa mite vectoring competence for DWV. Boxplots show development of DWV-A (upper panels) and DWV-B (lower panels) infection in individual naïve pupae after 6 day exposure with the phoretic, “Phor”, or pupal, “Pupal”, mites. Non-exposed control pupae - “None”, placed immediately after field collection (“Field-collected”) or following 12-day passage as phoretic mites on caged adult workers in the June (A) and September (B) experiments. C – Analysis of virus levels in worker honey bees after 12 day incubation with phoretic mites (“Phor”), pupal mites (“pupal”) mites, or without mites (“None”). Dots represent virus levels in individual pupae. Red letters above boxes indicate significantly and non-significantly different groups (pairwise t-test), NS – non-significant.

Such a dramatic effect of Varroa on the adult worker bee survival could be a result of either DWV-A or DWV-B infections. The RT-qPCR analysis of virus loads in adult bees of the September experiment at the day 12 showed significantly higher levels of both DWV-A and DWV-B in the mite-exposed cages (**Figure 3C**, “Phor”, “Pupal”) compared to the mite-free control cage (**Figure 3C**, “None”), *P* < 0.01. There were no significant differences between the levels of DWV-A or DWV-B in “Phor” and “Pupal” cages (**Figure 3C**) despite the difference in adult worker bee mortality. The caged mites also suffered high losses. In the June experiment, the number of live mites decreased from 67 to 18 for the phoretic mites, and from 50 to 3 for the pupal mites. In the September experiment the number of live mites in phoretic cage went from 60 to 19, and from 48 to 19 in the pupal cage.

All mites which survived for 12 days with caged adult bees were moved to naïve pupae for vector competence testing (**Figure 3A, B**, “12 day as phoretic”). After 6 days of mite feeding, we analyzed viral loads in the recipient pupa of the following groups: control with no mites (“None”), exposed to the mites of phoretic (“Phor”) and pupal (“Pupal”) origin (**Figure 3A, B**). We observed significantly higher levels of DWV-A and DWV-B for the mites which had a 12-day phoretic stage in cages (**Figure 3A, B**, “12 days as phoretic”) compared to those for the mites analyzed directly from the field (**Figure 3 A, B**,“Field-collected”), *P* <0.01 by t-test.

We also stratified mites according to their ability to cause overt-level virus infection in the recipient pupae. It was found that only 39.2 % (95% CI: 30.0 – 49.1%) of June and September field-collected mites (n = 97), were able to induce overt-level infection of DWV (DWV-A or DWV-B) in the pupae on which they were fed. After the 12-day phoretic passage on adult worker bees in cages, the infectiveness of surviving mites (n = 59) showed a significant increase, with 89.8% (95% CI: 79.2 – 95.6%) being able to cause high-level DWV infection (*P* < 0.0001 by chi-square test; **Figure 3A, B**).

After the phoretic cage passage, the ability to transmit both types of DWV was nearly equal, reaching 84.8% (95% CI: 73.3 – 92.0%) for DWV-A, and 86.4% (95% CI: 75.2 – 93.2%) for DWV-B (**Figure 3A, B**). These were significant increases compared to the direct infectiveness of the field mites, 23.7 % (95% CI: 16.3 – 33.1%) and 20.6 % (95% CI: 13.7 – 29.8%) for DWV-A and DWV-B, respectively.

### Varroa mites vectoring DWV-B had reduced lifespan

We assessed survival of each mite involved in vectoring competence tests by passaging mites to naïve white-eye pupae every six days until their death (**Figure 1**). We observed no seasonal trend in mite survival depending on the month of collection, average survival times (95% CI) for the mites of May, June, August, September, and October collections were 20.1 (95% CI: 16.7 - 23.4), 14.3 (95% CI:11.0 - 17.5), 24.3 (95% CI 20.5 - 28.2), 20.9 (95% CI 18.2 - 23.6), and 22.2 (95% CI 17.9 - 26.5) days respectively. By analyzing mites collected over 6 months (n = 256), no significant difference in survival was found between the field collected pupal and phoretic mites (*P* = 0.4, Pearson). We also investigated possible connections between the DWV type transmitted to the recipient pupae (**Figure 1**) and mite survival. We found that while there was no effect of transmission efficiency of DWV-A on mite survival (R = 0.063, p = 0.31, not significant) (**Figure 4A**), there was a significant negative correlation between the levels of DWV-B infection after mite feeding and mite survival, R = −0.14, *P* = 0.024 (**Figure 4B**). An even higher significance level was observed for the negative correlation between log-transformed ratios between DWV-B and DWV-A and mite survival time (R = −0.16, *P* = 0.0087) (**Figure 4C**). We stratified field-collected Varroa mites according to their ability to cause overt level of DWV-A or DWV-B in the recipient pupae after 6 days of feeding and analyzed the time mites survived after collection and placing on the recipient pupae (**Figure 4D, E**).

**Figure 4.**
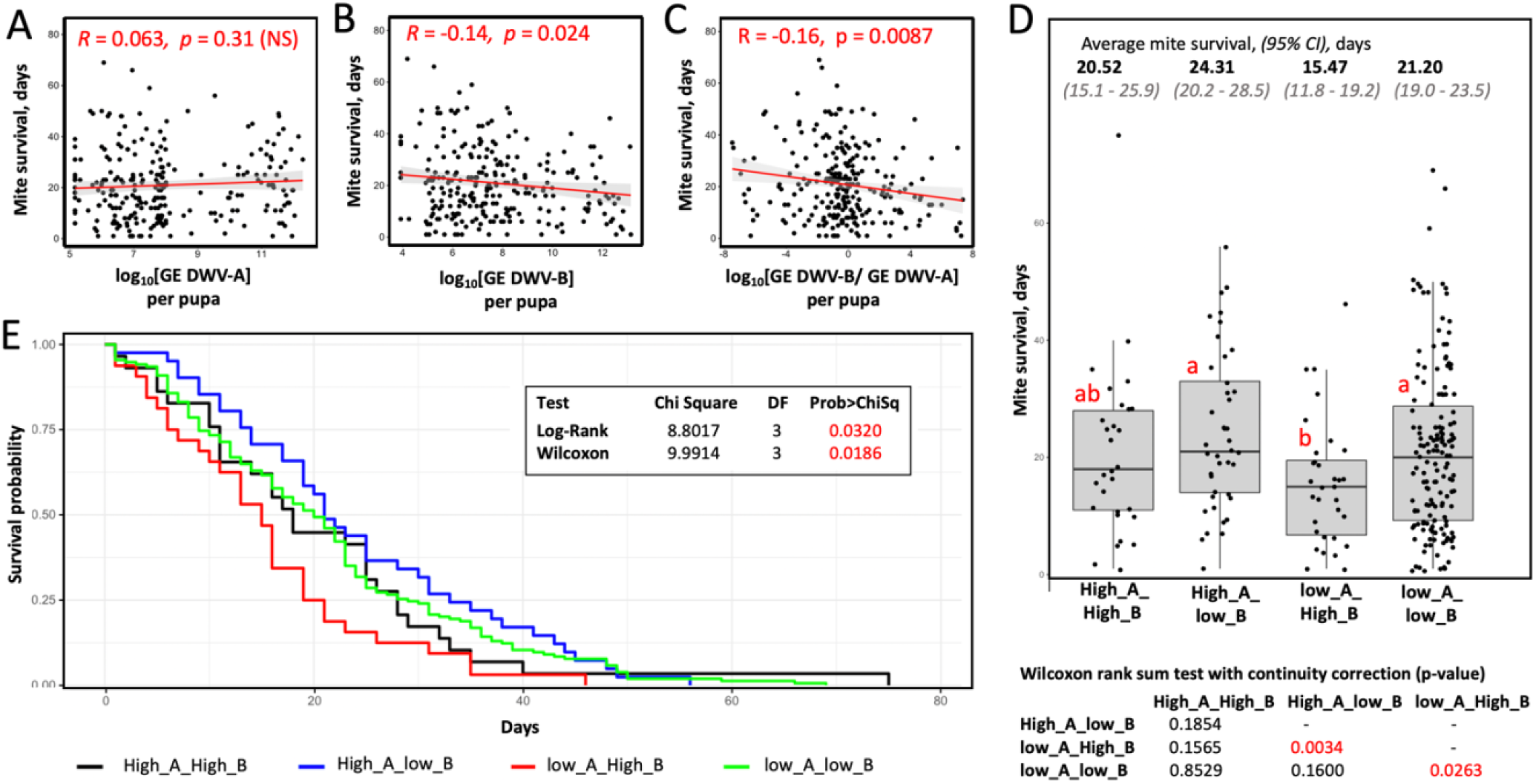
Connection between virus vectoring competence and survival time for field-collected mites. Shown are scatterplots, correlation values, and significance of correlation between the (A) levels of DWV-A, (B) levels of DWV-B, (C) log-transformed DWV-A to DWV-B ratio in the recipient pupae and the mite survival after filed collection. D – boxplot showing survival of the mites stratified according to their vectoring competence for DWV-A and DWV-B, threshold 10^9^ GE. Dots represent mite survival from the day of field collection. Red letters above boxes indicate significantly and non-significantly different groups (Wilcoxon test), the table summarizing results of Wilcoxon test of mite survival is below the graph. E-Survival probability analysis.

We found that mites, which induced overt DWV-B but not DWV-A infection after feeding on pupae, (Group “low_A_High_B”; n = 32) had the shortest survival time (15.5 days on average; 95% CI: 11.8 - 19.2 days), which was significantly shorter than that of the mites which induced overt DWV-A but not DWV-B infection (Group “High_A_low_B”; n = 41) with an average survival of 24.3 days (95% CI: 20.2 - 28.5), *P* = 0.0034, Wilcoxon test. The field mites which did not induce high levels of DWV-A or DWV-B, Group “low_A_low_B”, (n = 154) had an average survival of 21.2 days (95% CI: 19.0 - 23.5 days), which was not significantly different the Group “High_A_low_B”, but was significantly longer than in the case of the Group “low_A_High_B”, p = 0.026, Wilcoxon test. The mites which transmitted high levels of both DWV-A and DWV-B, Group “High_A_High_B” (n = 29) had an average survival of 20.5 days (95% CI: 15.1 - 25.9 days), which was not significantly different from other three groups (**Figure 4D, E**). Notably, “low_A_High_B” mites were found in all five field collection events, indicating that it could not be explained by seasonal influence. For example, there were 4 of 42 “low_A_High_B” mites in the June collection, which had shortest average survival of 14.3 days, 6 of 73 of such mites in the August collection (average survival of 24.3 days), and 9 of 50 for October (average survival of 22.2 days).

The mites which were transferred from cages (n=59) survived on average 8.72 days (95% CI: 7.1 - 10.3 days) after being placed on the vectoring competence test pupae (**Figure 1A, B**). Most of these mites (n = 48) induced high levels of both DWV-A and DWV-B in the pupa of and survived on average for 8.9 days (95% CI: 7.2 - 10.6 days). Reduced survival time of the mites following a 12-day phoretic stage compared to field mites could be a result of acquisition of DWV-B from the infected adult bees, differences in nutritional quality between adult workers and pupa, the additional 12 days of age, or a combination of all these possibilities. Therefore, we did not include the cage experiments mites to the survival analyses.

## Discussion

Vector-pathogen interactions may influence efficiency of vector-mediated transmission, as well as the spread and survival of the vectors themselves (26). Understanding these interactions is essential for development of disease control strategies and predictive modelling of pathogen spread. *Varroa destructor* and the viruses transmitted by this mite pose a major threat to apiculture (23), but there are many gaps in understanding the tri-partite interactions between honey bees, Varroa and viruses, including DWV. In this study we carried out quantitative assessment of vectoring competence of Varroa mites for honeybee viruses and measured the effects of virus vectoring on both bees and the mites themselves. This included direct tests of the mites’ ability to transmit DWV in controlled laboratory conditions in which individual mites were feeding on naïve honey bee pupae. We specifically tested infectiousness of Varroa mites in biological tests rather than quantifying loads of DWV in the mites because it was previously shown that a significant proportion of the virus load detected in mites by qRT-PCR may not be infectious (17), and therefore could not be an indicator of the ability of mites to transmit virus to recipient pupae during feeding.

We tested infectiousness of 256 field-collected mites, both phoretic and pupa-associated, which were directly collected from the same set of colonies on five occasions from May to October. Surprisingly, less than a half of these mites (39.8%) were able to induce overt-levels DWV-A or DWV-B infections in naïve pupae (**Figure 2**). Similar levels of infectiousness were found for phoretic and pupa-associated mites, and no obvious seasonal trends of infectiousness were observed (**Figure 2**). The overall ranges of DWV-A and DWV-B transmission were similar, but with significant variation between individual collection events (**Figure 2**). The possibility that such a low efficiency of DWV transmission from field-collected mites to pupae could be due to the pupal resistance to virus infection could be ruled out, because a significantly higher DWV transmission rate (89.8 %) was observed in pupae collected from the same colonies after 12 days with caged adult worker bees as phoretic (**Figure 3A, B**).

The experiments that maintained the mites on adult worker bees (**Figure 1**) were designed to investigate the effect of extended phoretic stage on the ability of the mites to transmit viruses. We observed significantly higher levels of DWV-A and DWV-B in caged worker bees exposed to mites as opposed to mite-free controls after 12 days (**Figure 3C**), indicating that bees were infected by introduced mites. A trophallactic transmission of viruses between bees also could contribute to the spread of infection (25, 27). As a result, the mites which had a low vectoring competence for DWV at the time of their collection from the field colonies could acquire DWV by feeding on the infected bees in cages. The cage experiments also clearly showed that Varroa infectiousness could be increased during broodless periods and surviving mites might have a higher potential to transmit the virus. In our cage experiments, almost all mites became highly infective following incubation on adult bees, suggesting that interruptions in honey bee brood production (i.e. ‘brood breaks’) which decrease Varroa infestation (24), might be less effective in decreasing DWV loads because mites which survived brood interruptions could be highly infectious and waiting to transmit the virus to pupae. Our cage experiments findings might explain the results of recent colony study (28), which showed that 20 day-long queen caging resulting in the absence of brood and Varroa fall negatively impacted on colony strength and survival. This highlights the need to consider impact of prolonged phoretic mite stage on circulation of DWV.

A high proportion of field-collected Varroa which were non-infectious in pupal tests (**Figure 2A**) but were quick to increase infectiousness following phoretic exposure to bees (**Figure 3A, B**). This observation agrees with the model of non-replicative and, possibly, non-persistent vectoring of DWV-A (17), but apparently contradicts evidence for widespread replication of DWV-B in Varroa (16, 29), which should result in the maintenance of vector competence for DWV-B long-term. We did not observe an increase of vectoring competence for DWV-B in field mites during the season, despite obvious presence of DWV-B. However, this disagreement could be explained by the results of the mite survival analysis (**Figure 4**), which showed different effects of DWV-A and DWV-B on the time mites remained alive on pupae in capsules (**Figure 1**). We found that the mites which were able to transmit only DWV-B to the recipient pupae had a significantly reduced lifespan compared to the mites which efficiently vectored DWV-A or did not cause high levels of either DWV-A of DWV-B (**Figure 4**). These results also excluded a possibility that the increase of infectiousness after 12-day phoretic stage on adult worker bees in cages was a consequence of better survival of mites that were able to cause DWV-A or DWV-B infections compared to mites with low vectoring potential.

The negative impact of DWV-B but not DWV-A vectoring on Varroa survival is likely a consequence of the ability of DWV-B, but not DWV-A, to replicate in the mites (16, 29). It has been reported that viral infections have negative effects of fitness, reproduction, and survival of their mosquito vectors (30, 31). DWV-B, which was associated with increased overwintering losses of honeybees (32), has become more widely widespread than DWV-A at least in the USA (33), but complete elimination and replacement of DWV-A by DWV-B has not been observed. We propose that reduced survival of Varroa carrying DWV-B may provide negative feedback to the dominance of DWV-B in honey bee colonies and could explain why DWV-B has not completely replaced DWV-A.

It is very likely that reduction of DWV-B loads during the winter broodless season, as reported by Traynor et al. (34), may be a result of increased mortality of both the virus-infected adult bees and the vectoring mites. Further systematic large-scale assessment of DWV-B loads and Varroa infestation levels throughout the season are required to determine the effect of mutual influence of DWV-B and Varroa.

The study of interactions between Varroa and DWV types A and B may provide an insight into general understanding on evolution of vector-pathogen interactions. A pathogen may maximize vector-mediated transmission rate by evolving an ability to replicate in its vectors, but this might have a negative impact on vector fitness and survival (4, 30). Therefore, both replicative and non-replicative transmission modes have their advantages. DWV was present in honey bees before the arrival of Varroa (11), and it is likely that differences between DWV-A and DWV-B in their abilities to replicate in Varroa are recent adaptations. Therefore, further comparison of DWV-A and DWV-B Varroa-mediated circulation may help to understand the dynamics of pathogen vectoring in the face of reduced vector fitness. Information on vectoring competence of Varroa and the factors influencing it, as well the effects of honey bee viruses on vectors and their hosts, will unite Varroa and virus dynamics in honey bee colonies and help resolve their impacts on colony disease and survival.

## Author Contributions

EVR, FPF and SCC had contributed equally to this work and share senior authorship. EVR, FPF and SCC: conceptualization, project administration. EVR, SCC, YC - funding acquisition. FPF, YC, ZSC: resources. EVR, FPF, JDE, SCC, ZSL: methodology. FPF, EVR, CR and SCC: investigation and formal analysis. EVR, FPF and SSC original draft preparation. EVR, FPF, SSC, YC, JDE writing. SSC, EVR: supervision. All authors have read and agreed to the published version of the manuscript.

